# Assessment of segregation variance estimates from derivation, simulations, and empirical data in autotetraploid species exemplified in potato

**DOI:** 10.1101/2025.08.13.670171

**Authors:** Po-Ya Wu, Kathrin Thelen, Stefanie Hartje, Katja Muders, Vanessa Prigge, Benjamin Stich, Delphine Van Inghelandt

## Abstract

The optimal choice of parents and crosses and, therefore, the prediction of the segregation variance are of high relevance to maximize genetic gain in breeding programs. Several methods have been developed for the prediction of segregation variance, including correlation with genotypic diversity, progeny simulations, or algebraic derivations in case of a diploid inheritance. To the best of our knowledge, no algebraic derivation using parental genotypic information is available to predict segregation variance for autotetraploid species. The objectives of our study were to (1) derive algebraic derivation based on linkage disequilibrium (LD) between linked loci to predict the segregation variance in autotetraploid species; (2) compare the performance of segregation variance estimated based on simulated progenies and the algebraic derivation; (3) investigate by simulations how experimental parameters affect the accuracy of segregation variance prediction; and (4) compare the segregation variance estimated in empirical data of potato and the one based on the algebraic derivation. The segregation variance estimated by the developed derivations showed very high correlations with the one observed in large simulated progenies, but those were lower when phased parental haplotypes were not available or family size decreased. The correlation between segregation variance estimated by the developed derivation and the empirical data was low. This could be attributed to the small family size used in the study, which we could show to increase LD between unlinked loci. The proposed algebraic derivations promise to be a precise alternative to simulations to help breeders in optimizing their family choices and sizes considering the segregation variance.

## INTRODUCTION

The breeders’ goal is to develop new varieties with increased performance in comparison to the existing ones. In the narrow sense, this means to create segregating populations via crosses among potential parents, and perform selection in the progenies to increase their performance in comparison to those of their parents. This increase in performance between the generations before and after selection is known as the genetic gain (ΔG) (Fisher, 1918). Lush (1937) developed the formula to quantify the expected genetic gain ΔG = H^2^S (where H^2^ is the heritability and S the selection differential), and this equation is commonly known as the breeder’s equation. Later, Falconer and Mackay (1996) quantified the expected genetic gain as ΔG = iHσ_G_, where i is the selection intensity, H the square root of heritability, and σ_G_ the genetic standard deviation. As the number of families that can be established and considered in a breeding program each year among all the potential parents is almost infinite, it has to be restricted due to time and resources constraints. Therefore, breeders have to choose between all possible crosses. To help the breeders in their choices, Schnell and Utz (1975) proposed the usefulness of a cross for selection (UC), a parameter combining the expected progeny mean µ and the genetic gain possible to realize within this cross: UC = µ + iHσ_G_. From this formula, it becomes obvious that not only the progeny mean but also the segregation variance is important in the breeding process, i.e., from families with a similar high progeny mean, the one with a higher segregation variance should be preferred (Neyhart and Smith, 2019). Thus, developing approaches to quantify the segregation variance is pivotal to optimize choices in a breeding program.

Several studies have investigated the prediction of segregation variance using phenotype distance (Utz et al., 2001), pedigree, genetic distance, however, without success, as reviewed in Mohammadi et al. (2015) and Lehermeier et al. (2017b). With the development of dense genome-wide markers and the advent of genomic prediction (GP) models, the marker effects can be well estimated (Meuwissen et al., 2001). Therefore, the segregation variance can now be estimated either via simulated progenies (Bernardo, 2014), or via algebraic development (e.g., Bonk et al., 2016; Lehermeier et al., 2017b; Osthushenrich et al., 2017; Allier et al., 2019; Wolfe et al., 2021; Niehoff et al., 2024) incorporating the estimated marker effects from the GP model. To achieve the former approach, several simulation tools used to generate the progenies, such as PopVar (Mohammadi et al., 2015) or AlphaSimR (Gaynor et al., 2021), have been developed. Some of these tools can be applied for a wide range of both diploid and autotetraploid species. However, these simulators require key inputs such as genetic map information and phased parental haplotypes. A comprehensive summary of all simulation tools and their functionalities and assumptions is available from Stich et al. (2025, under review).

Using simulated progenies to estimate segregation variance can, however, be computationally intensive, especially when a large number of breeding crosses needs to be evaluated, which is typical for breeding programs (Lehermeier et al., 2017b). An alternative to estimate segregation variance is based on algebraic derivations. Several derivations considering linkage disequilibrium (LD) between linked loci, recombination rate, as well as phased parental haplotypes have been derived recently to predict the segregation variance (Bonk et al., 2016; Lehermeier et al., 2017b; Osthushenrich et al., 2017; Allier et al., 2019; Wolfe et al., 2021; Niehoff et al., 2024). However, these derivations are developed based on diploid inheritance and are not suitable for other important crops with autopolyploid genomes, such as yams, sugarcane, and potato.

Another concern in autopolyploid species is that inferring the phased parental haplotypes is in practice a costly and time-consuming process due to the complexity of their genome (e.g., Sun et al. 2022, Sun et al. 2025). For such cases, the segregation variance estimation based on simulated progenies is not applicable due to the absence of phased parental haplotypes. In contrast, estimating segregation variance with unknown phased parental haplotypes could be done via algebraic derivation under the assumption of the absence of LD between linked loci. Therefore, investigating the possibility to estimate segregation variance via algebraic derivation when phased parental haplotypes are unknown is of interest.

A further aspect about algebraic derivation is that they are developed under the assumption of a population in equilibrium, that is, infinite family size, which could theoretically yield more precise segregation variance estimation than simulated approaches. This aspect needs to be assessed. Furthermore, in practical breeding programs, limited budget, and resources can restrict the allocations of family size and number of families in field trials or greenhouses, thereby affecting the selection of potential crosses. Furthermore, reproductive issues leads to uneven family size across different families. Therefore, the variability of number of families, the family size, and its distribution across the assessed families could prevent to reveal the true genetic variance of a trait for each family. The impacts of the above-mentioned factors on segregation variance estimation need to be investigated, which can be studied by computer simulations.

In this study, we compared several methods to assess the genetic variance of F1 segregating populations in autotetraploid species using potato as example. Our objectives were to: (1) derive algebraic derivation based on LD between linked loci to predict the segregation variance in autotetraploid species; (2) compare the per-formance of segregation variance estimated based on simulated progenies and the algebraic derivation; (3) investigate by simulations how experimental parameters affect the accuracy of segregation variance prediction; and (4) compare the segregation variance estimated in empirical data and the one based on the algebraic derivation.

## MATERIAL AND METHODS

In this study, we compared several methods – algebraic derivations, simulation, and empirical data – to estimate the segregation variance in autotetraploid species using potato as an example. The evaluation scheme and comparison framework are shown in Figure 1.

**Figure 1:**
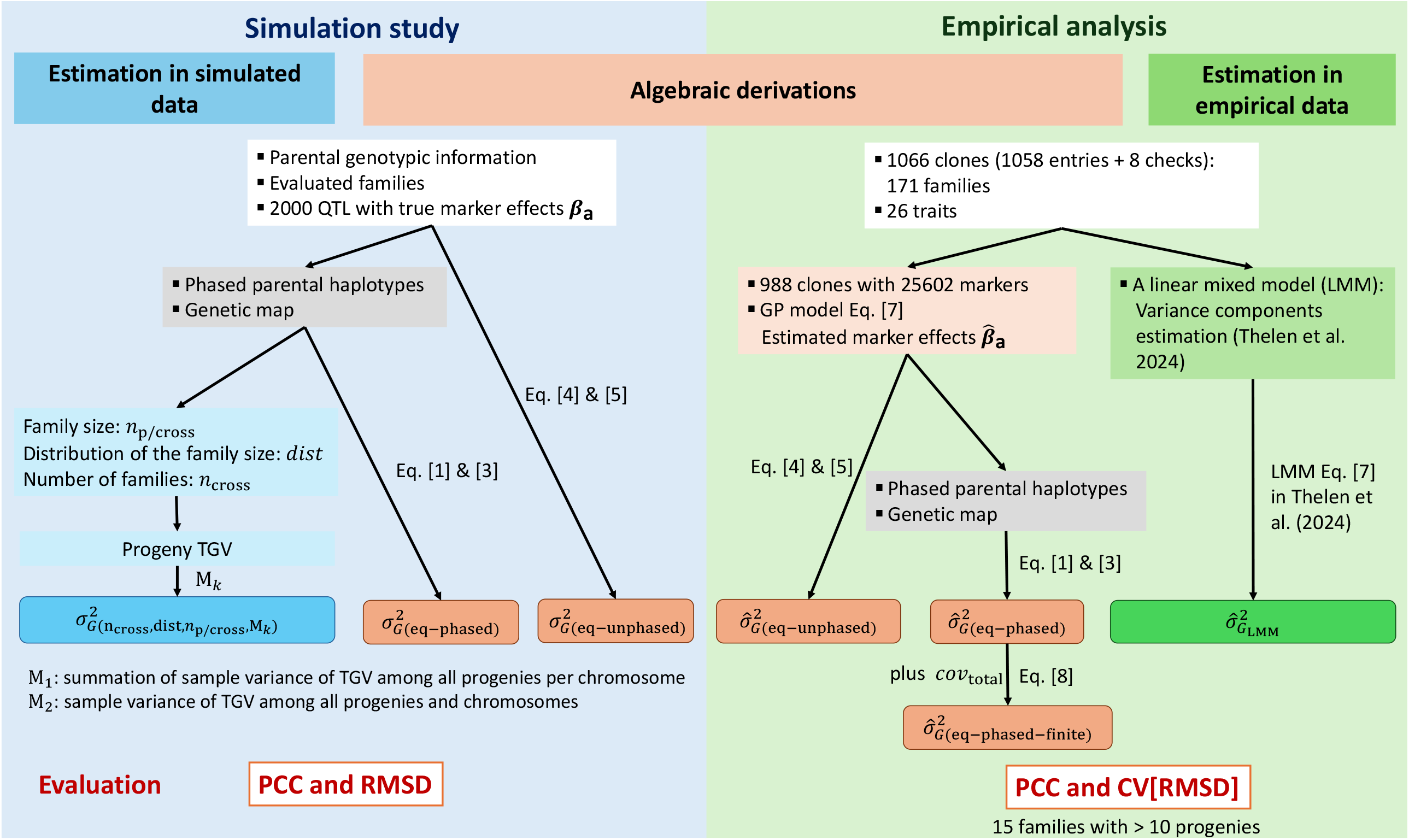
Scheme of the evaluation of segregation variance estimation based on simulation, algebraic derivations, and empirical data. PCC: Pearson correlation coefficient; RMSD: root mean square deviation; CV[RMSD]: the coefficient of variation of the RMSD; and TGV: true genetic values.

### Derivation of the additive segregation variance

We focused on developing an algebraic derivation to estimate the segregation variance considering bi-allelic QTL, given known estimated additive marker effects - β_a_, excluding non-additive effects, and assuming no double reduction.

We assume two parents, P1 and P2, of a highly heterozygous and autotetraploid crop, and their known genotypic information for p biallelic molecular markers being X_P1_ and X_P2_, which are two 1 × p matrices. The genotypic information involves five genotype classes, coded as 0, 1, 2, 3, and 4 and the additive marker effects are defined as β_a_, which is a p × 1 vector. In heterozygous species, the members of a F1 family established from the P1 × P2 cross are segregating. Such a family is designated in the following as segregating population. In case of partial homozygosity, concerned loci will not segregate in F1, however, the algebraic derivation is still valid. The genotypic information of the n F1 progenies can be denoted as X_P1×P2_, which is a n × p matrix. The additive segregation variance can be derived as:

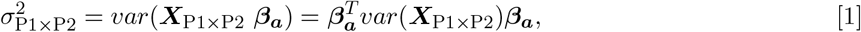

where 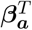 is the transpose of β_a_, and var(X_P1×P2_) the additive genotypic variance of the progenies, which is a p × p matrix. Each non-diagonal element in var(X_P1×P2_) is the covariance between two loci (A and B), noted as cov(x_A_, x_B_). Each diagonal element in var(X_P1×P2_) is the variance of the locus itself, noted as var(x_A_). Until here and except for the kind of genotypic information used (five genotype classes instead of three), the derivation is similar to the one for diploid species. In the following, formulae were derived for two scenario: (i) the phased parental haplotypes are known, and (ii) the phased parental haplotypes are unkown.

### With phased parental haplotypes

In a first scenario, we assumed that phased parental haplotypes and the recombination rate (c) between two loci A and B are known. Therefore, cov(x_A_, x_B_) can be derived based on the LD between pairs of loci among all possible combinations of genotypes produced by a set of two parents. Here, we assume only random bivalents are formed during meiosis, implying that no double reduction occurs (Gallais, 2003). Thus, the segregation at two linked loci allows the formation of 36 different gametes for each genotype (Suppl. Figure S1 revised from Figure 1.4 in Gallais 2003). Each gamete has a probability as given in Table 1. We denote that a genotype at two linked loci, A and B, can be expressed as A_1_B_1_/A_2_B_2_/A_3_B_3_/A_4_B_4_ (see Suppl. Figure S1). Only bi-allelic loci are considered in our study, and, thus, any A_h_ and B_h_ is coded as 0 or 1 for representing the reference allele or the alternative one, where h indicates the 1,2,3, or 4^th^ homolog.

**Table 1:** The probabilities of 36 gametes from a genotype, A_1_B_1_/A_2_B_2_/A_3_B_3_/A_4_B_4_, in a scenario with two loci A and B, and bivalent formation during meiosis. *c* is the recombination rate between loci A and B.

|  | Gamete | Type* | Probability |
| --- | --- | --- | --- |
| 1 | $A_1B_1/A_2B_2$ | 2P | $\frac{(1-c)^2}{6}$ |
| 2 | $A_1B_1/A_3B_3$ | 2P | $\frac{(1-c)^2}{6}$ |
| 3 | $A_1B_1/A_4B_4$ | 2P | $\frac{(1-c)^2}{6}$ |
| 4 | $A_2B_2/A_3B_3$ | 2P | $\frac{(1-c)^2}{6}$ |
| 5 | $A_2B_2/A_4B_4$ | 2P | $\frac{(1-c)^2}{6}$ |
| 6 | $A_3B_3/A_4B_4$ | 2P | $\frac{(1-c)^2}{6}$ |
| 7 | $A_1B_2/A_3B_4$ | 2R | $\frac{c^2}{6}$ |
| 8 | $A_1B_2/A_4B_3$ | 2R | $\frac{c^2}{6}$ |
| 9 | $A_2B_1/A_3B_4$ | 2R | $\frac{c^2}{6}$ |
| 10 | $A_2B_1/A_4B_3$ | 2R | $\frac{c^2}{6}$ |
| 11 | $A_3B_1/A_4B_2$ | 2R | $\frac{c^2}{6}$ |
| 12 | $A_1B_3/A_2B_4$ | 2R | $\frac{c^2}{6}$ |
| 13 | $A_1B_1/A_3B_4$ | 1P1R | $\frac{c(1-c)}{12}$ |
| 14 | $A_1B_1/A_4B_3$ | 1P1R | $\frac{c(1-c)}{12}$ |
| 15 | $A_2B_2/A_3B_4$ | 1P1R | $\frac{c(1-c)}{12}$ |
| 16 | $A_2B_2/A_4B_3$ | 1P1R | $\frac{c(1-c)}{12}$ |
| 17 | $A_3B_3/A_1B_2$ | 1P1R | $\frac{c(1-c)}{12}$ |
| 18 | $A_4B_4/A_1B_2$ | 1P1R | $\frac{c(1-c)}{12}$ |
| 19 | $A_3B_3/A_2B_1$ | 1P1R | $\frac{c(1-c)}{12}$ |
| 20 | $A_4B_4/A_2B_1$ | 1P1R | $\frac{c(1-c)}{12}$ |
| 21 | $A_1B_1/A_2B_4$ | 1P1R | $\frac{c(1-c)}{12}$ |
| 22 | $A_1B_1/A_4B_2$ | 1P1R | $\frac{c(1-c)}{12}$ |
| 23 | $A_3B_3/A_2B_4$ | 1P1R | $\frac{c(1-c)}{12}$ |
| 24 | $A_3B_3/A_4B_2$ | 1P1R | $\frac{c(1-c)}{12}$ |
| 25 | $A_2B_2/A_1B_3$ | 1P1R | $\frac{c(1-c)}{12}$ |
| 26 | $A_4B_4/A_1B_3$ | 1P1R | $\frac{c(1-c)}{12}$ |
| 27 | $A_2B_2/A_3B_1$ | 1P1R | $\frac{c(1-c)}{12}$ |
| 28 | $A_4B_4/A_3B_1$ | 1P1R | $\frac{c(1-c)}{12}$ |
| 29 | $A_1B_1/A_2B_3$ | 1P1R | $\frac{c(1-c)}{12}$ |
| 30 | $A_1B_1/A_3B_2$ | 1P1R | $\frac{c(1-c)}{12}$ |
| 31 | $A_4B_4/A_2B_3$ | 1P1R | $\frac{c(1-c)}{12}$ |
| 32 | $A_4B_4/A_3B_2$ | 1P1R | $\frac{c(1-c)}{12}$ |
| 33 | $A_2B_2/A_1B_4$ | 1P1R | $\frac{c(1-c)}{12}$ |
| 34 | $A_3B_3/A_1B_4$ | 1P1R | $\frac{c(1-c)}{12}$ |
| 35 | $A_2B_2/A_4B_1$ | 1P1R | $\frac{c(1-c)}{12}$ |
| 36 | $A_3B_3/A_4B_1$ | 1P1R | $\frac{c(1-c)}{12}$ |

\*: 2P – with parental chromosome association, i.e., without any recombination; PR – with one parental chromosome association and one recombinant chromatid; and 2R – with two recombinant chromatids.

When two parents are crossed, 1296 (= 36 × 36) genotypes are possible at two loci in their F1 progenies. All possible genotypes and their probabilities are shown in Table 2 and Suppl. Table S1. The covariance between the two loci can be expressed by:

**Table 2:**
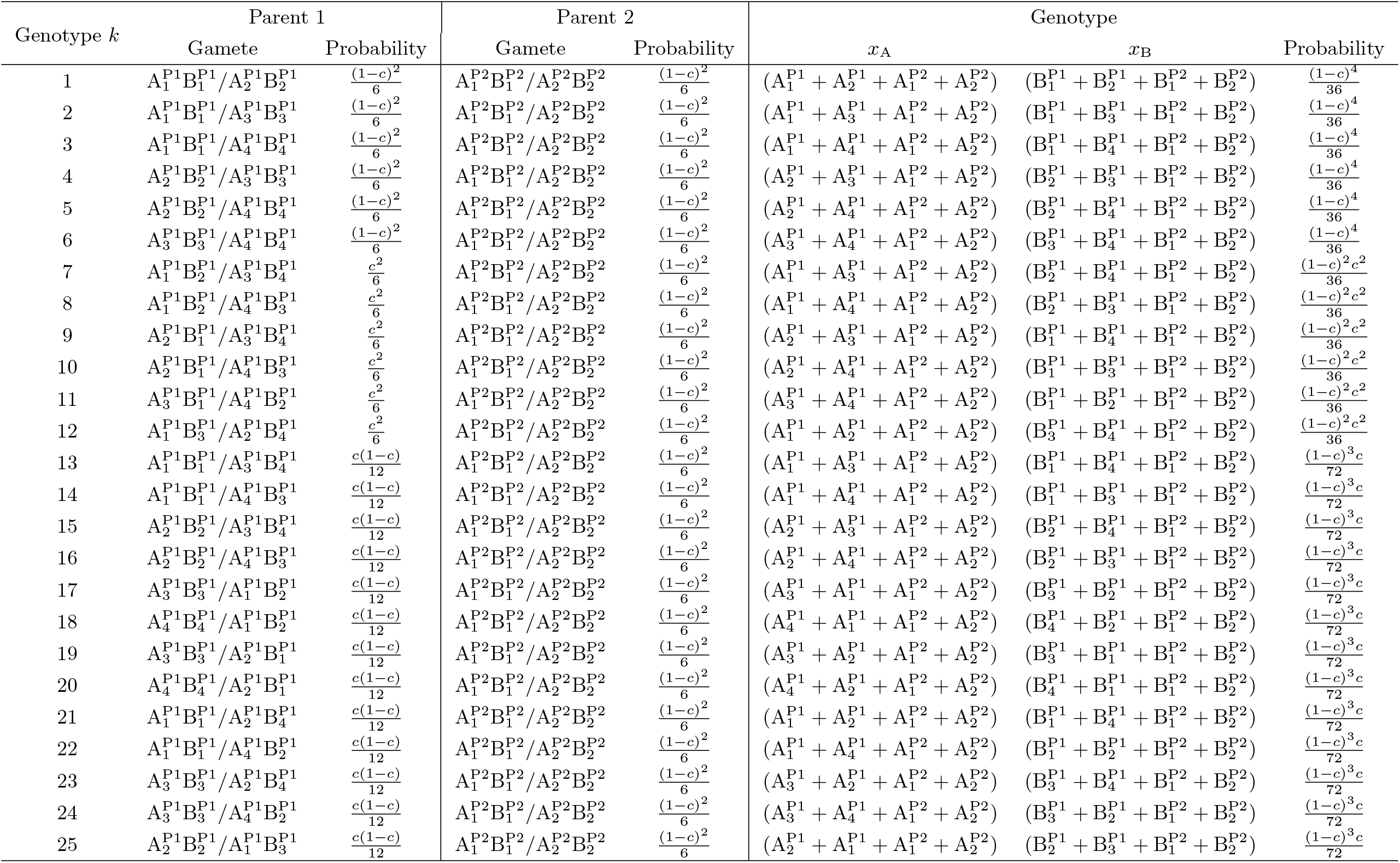
The 1296 possible genotypes in a F1 family of a bi-parental cross and their corresponding probabilities, depending on the parental gametes and their probability. *c* is the recombination rate between the two loci A and B. Here only the first 25 genotypes are shown. Information for all 1296 genotypes is shown in Suppl. Table S1.

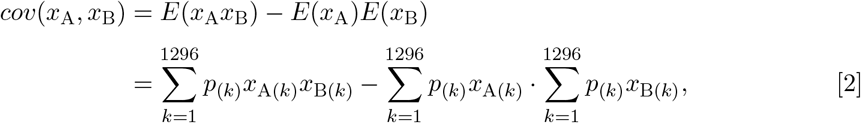

where p_(k)_ is the probability of the k^th^ genotype from Table 2 (last column), and x_A(k)_ (x_B(k)_) the genotypic indicator contributed by the two parental gametes for locus A (B) at the k^th^ genotype, corresponding to the 6^th^ (7^th^) column in Table 2. In our study, only additive effects are considered, and therefore equation [2] can be shortened to:

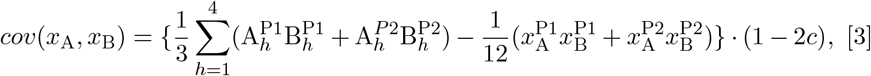

where 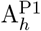 and 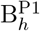 (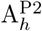and 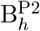) denote the genotypic indicator (0/1 for the refer-ence/alternative allele, respectively) at the h^th^ homolog of A and B loci for P1 (P2), and 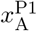 and 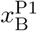 (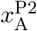 and 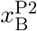) the genotypic indicator of A and B loci for P1 (P2) ranging from 0 to 4, where 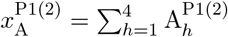 and 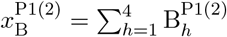.

In equation [3], the covariance is 0 if c = 0.5, that is, the two loci are located on different chromosomes. Furthermore, the additive variance of a locus itself can be simplified from equation [3], where c is set to 0 and locus B replaced by A, and written as:

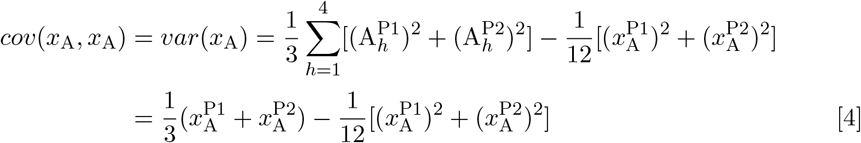

### Without phased parental haplotypes

The derivation of additive segregation variance 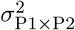 (Eq. [1]) can be simplified when phased parental haplotypes are unknown. In this case, the LD between two loci cannot be taken into account. That is, only the additive variance for each locus itself can be estimated, with the off-diagonal elements of var(X_P1×P2_) being 0. Subsequently, the additive segregation variance can be expressed in that case as:

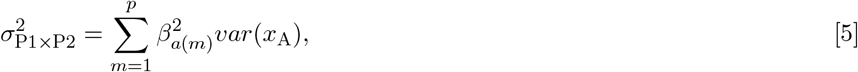

where var(x_A_) was equal to equation [4].

### Simulation study

To validate the accuracy of the derived formulae, we first compared the prediction accuracy of segregation variance estimation based on equation [1], [3], and [5] and the variance estimated by simulated progenies. Further, to investigate how different experimental settings affect the performance of segregation variance prediction, several parameters, including (i) the family size per segregating population, (ii) the distribution of the family sizes across segregating populations, and (iii) the number of families in a breeding program, were varied in our simulations.

### Parameters of the simulation

For the simulations, we used 100 autotetraploid potato clones from a resequencing panel (Baig et al. in preparation) as parents of a half-diallel generating 4950 crosses. For the 100 parents, 49,125 markers across 12 chromosomes were available after quality control (Wu et al., 2025), and their genetic map information was estimated as described in Wu et al. (2023). Here, the phased haplotypes of these markers were randomly assigned. The genetic map distance (d, Morgan) between two linked loci was converted to a recombination rate (c) using the Haldane mapping function (Haldane, 1919): c = 0.5(1 − e^−2d^). A random set of 2000 markers was considered as QTLs for an index trait summarizing all target traits of potato breeding. True additive QTL effects β_a_ were drawn from a gamma distribution with k = 2 and θ = 0.2, where k and θ are shape and scale parameters, respectively. The true genotypic value for each genotype (g) was then defined as g = x_a_β_a_, where x_a_ is the additive genotypic indicator across all QTLs with a dimension of 1 × 2000. As only additive effect for each QTL were considered, the true genotypic value was identical to the breeding value.

Different family sizes (n_p/cross_) could affect the precision of segregation variance estimates using simulated progenies. To investigate this aspect following a realistic scenario for breeders but also assess which family size would be needed to assess perfectly segregation variance, we let n_p/cross_ vary from 10, 20, 50, 100, 200, 500, 1000, 2000, to 5000. Furthermore, in practical potato breeding programs, the distribution of the family sizes at A clone stage can follow different distributions (Suppl. Figure S2). Thus, we considered in our simulations the distribution of the family sizes across segregating populations, denoted as distr, to follow either a uniform or a gamma distribution, U or Γ, respectively. The last parameter, the number of families (n_cross_), varied from 10, 50, 100, 250, to 500.

In total, there were 90 different experimental settings, which were called scenarios hereafter.

### Segregation variance estimation

In each scenario, the segregation variance was estimated by (1) the summation of sample variance of true genetic values among all simulated progenies per chromosome, thus ignoring the covariance between QTLs on different chromosomes, denoted as method 1 (M_1_); and (2) the sample variance of true genetic values among all sim-ulated progenies and chromosomes, thus considering the covariance between QTLs on different chromosomes, denoted as Method 2 (M_2_). The estimated variance from simulated progenies was called 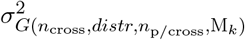 hereafter, where M_k_ represents Method k.

Furthermore, the expected segregation variance was estimated by:

- the equations [1] & [3] with phased parental haplotypes, true marker effect β_a_

and called 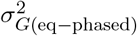 hereafter.

- the equations [4] & [5] with unphased parental haplotypes, true marker effect

β_a_, called 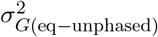 hereafter.

### Evaluation

We compared the segregation variance obtained from the two methods that were based on simulated progenies 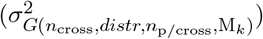 and the ones obtained by the derivations (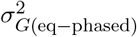 and 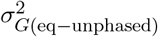) by calculating the Pearson correlation coefficient (PCC) and their root mean square deviation (RMSD) in each scenario.

In addition, the PCC and RMSD between 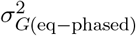 and 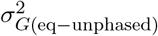 were estimated.

To avoid effects due to sampling in our stochastic simulations, all results were based on 30 independent simulation runs. The positions of 2000 QTLs were changed in each run, but their effect size was fixed across 30 runs. The median of PCC and RMSD were taken across the 30 runs. In this study, the simulated progenies were generated by AlphaSimR (Gaynor et al., 2021) with “quadProb = 0” in the simulation parameters, implying that no double reduction occurs in our simulated progenies.

### Assessment of the segregation variance in empirical data

To assess the application of our algebraic derivation in real breeding situations, we compared the segregation variances estimated by the derived formula using marker effects estimated in empirical potato populations and the empirical segregation variance estimated from a linear mixed model (LMM).

### Genetic materials and phenotypic characterization

A total of 1058 entries corresponding to the A clone stage which were a subset of the breeding material from the three breeding companies Europlant (EUROPLANT Innovation GmbH & Co. KG, with 300 entries), Norika (NORIKA GmbH, with 300 entries), and SaKa (SaKa Pflanzenzucht GmbH & Co. KG, with 458 entries) (Suppl. Table S2) were used in this study together with eight check varieties. The 1058 entries belonged to 171 full-sib families, which were designated in the following as segregating populations. The family size of one segregating population varied between one and 38 (Suppl. Figure S2).

The above-mentioned 1066 entries and checks were evaluated from 2019 to 2021 in field experiments in several locations in Germany, resulting in a total of 15 different environments (year × location combinations; Suppl. Table S2). Within the environments, the clones were organized in a block system with rows and columns following an augmented design. The eight checks were replicated eight times in each environment, while the entries were cultivated within each environment only once. The number of plants per plot ranged from nine to 20, depending on the respective environment (Suppl. Table S2).

Phenotypic data were recorded on an individual plot basis for 26 different traits (Table 3). Trait values were either assessed as a rating from 1 to 9, given in the form of a percentage value, or measured. More details about the experimental design, phenotypic data processing and analysis were described in Thelen et al. (2024), and the adjusted entry means calculated from the model with heterogeneous error variance (Thelen et al., 2025, in press) were used for the further analyses.

**Table 3:** Abbreviations and units for the evaluated traits considered in our study. The heritabilities on an entry mean basis (h^2^) were calculated for each trait as described by Thelen et al. (2024).

| Abbreviation | Trait | Unit | $h^2$ | Note |
| --- | --- | --- | --- | --- |
| CR4 * <sup>†</sup> | crisps color after storage at 4°C | 1-9 | 0.81 | 1 = bad quality (e.g. very dark), 9 = good quality |
| CR8 * | crisps color after storage at 8°C | 1-9 | 0.89 | 1 = bad quality (e.g. very dark), 9 = good quality |
| DEV | foliage development | 1-9 | 0.76 | 1 = not grown to very weak development, 9 = superior / extraordinary growth |
| DSC * | after cooking discoloration | 1-9 | 0.64 | 1 = very dark, 9 = no discoloration |
| EMR | emergence | 1-9 | 0.77 | rate of emergence from the soil, 1 = very poor, 9 = very good |
| EYE | eye depth | 1-9 | 0.86 | 1 = very deep, 9 = very flat |
| FLE | flesh color | 1-9 | 0.89 | 1 = white, 9 = blue/purple |
| FRI * | french fry color | 1-9 | 0.79 | 1 = bad quality (e.g. very dark), 9 = good quality |
| IMP | general impression | 1-9 | 0.69 | 1 = deficiencies, 9 = very good |
| MAT | maturity | 1-9 | 0.87 | 1 = still flowering, 9 = dead, relative to checks |
| PPO | polyphenol oxidase activity | 1-9 | 0.91 | tuber flesh after DL-DOPA incubation, 1 = no color change, 9 = very dark coloration |
| RHI <sup>†</sup> | rhizoctonia symptoms | 1-9 | 0.57 | 1 = >90 % infested, 9 = no symptoms |
| SCA | scab symptoms | 1-9 | 0.44 | 1 = >90 % infested, 9 = no symptoms |
| SHD | tuber shape diagonally | 1-9 | 0.53 | 1 = round, 9 = flat |
| SHL | tuber shape longitudinally | 1-9 | 0.92 | 1 = round, 9 = long |
| SIZ | tuber size | 1-9 | 0.81 | 1 = small, 9 = big |
| SKC | skin color | 1-9 | 0.88 | 1 = cream, 9 = blue/purple |
| TEX * | texture | 1-9 | 0.57 | 1 = tuber falls completely apart, 9 = tuber stays tightly together |
| TST * | taste | 1-9 | 0.57 | 1 = very strong deficits (e.g. bitter), 9 = nice potato taste |
| SKT | skin texture | 1-4 | 0.85 | 1 = plain, 4 = cracked |
| BRU * | susceptibility to bruising | % | 0.89 | 5 kg tuber sample |
| STA | starch content | % | 0.95 | measurement with automatic starch scale |
| TUL * | proportion of large tubers >65mm | % | 0.89 | weighting after grading |
| TUN * | proportion of normal tubers 35-65mm | % | 0.85 | weighting after grading |
| TUS * | proportion of small tubers <35mm | % | 0.87 | weighting after grading |
| YLD | total tuber yield | kg | 0.83 | normalized to a 16 plant plot |

\*: These traits were only assessed in years 2020 and 2021.
<sup>†</sup>: The traits CR4 and FRI were not evaluated for all clones, but only for those clones that belonged to the corresponding market segment, which was crisp production for CR4 and french fries production for FRI (see details in Thelen et al. (2024)).

### Genomic characterization

Out of the 1066 phenotyped clones, 988 (980 entries and 8 checks) have been genotyped using a SNP array with 224,009 high-quality SNP markers (Baig et al. in preparation). The 980 entries belonged to 163 full-sib families. The family size within one segregating population varied between one and 29 (Figure S2). In order to phase the parental haplotype robustly from progeny information, genotypic information of both parents is required, and the family size must be larger than 10. Thus, only 15 of 163 families were considered, and 26 parents were involved in their creation. We filtered out those markers that were not part of the dAg1 v1.0 reference genome (Freire et al., 2021), and the two parents for each cross with the remaining 206616 markers were phased using PolyOrigin (version v1.0.1; Zheng et al., 2021). Furthermore, during the parental phasing, the markers were removed if they did not fit into the marker sequence. Thus, the number of common phased markers across 15 families was 25602. In the last step, the physical map of the 25602 markers was converted into the genetic map using the Marey map constructed by Wu et al. (2023). From that in turn, the recombination rate c was estimated using the Haldane mapping function (Haldane, 1919).

### Estimation of empirical segregation variance

The LMM (see model [7] in Thelen et al. (2024)) was used to estimate the segregation variance for each family, which was called the empirical segregation variance. To do so, heterogeneous variances for the clones in each family were assumed in the model [7] of Thelen et al. (2024). Subsequently, the intra-family variance for each segregating population individually was estimated (Thelen et al., 2024) and denoted as 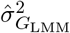 hereafter.

### Prediction of segregation variance with GP and derivations

In order to predict the segregation variance using the derived formula with or without the consideration of parental haplotype information in the empirical breeding population, i.e., 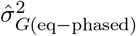 and 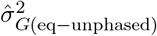, respectively, the following three components are required: (i) the phased parental haplotypes, (ii) the genetic map, and (iii) the estimated marker effects (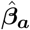instead of true marker effect). The first two components are available from the former section. The third component 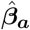 was obtained using a general GP model:

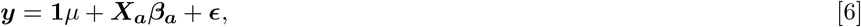

where y was the vector of the above-described adjusted entry means of each examined trait, 1 the unit vector, µ the general mean, X_a_ the matrix of the additive genotypic data information, β_a_ the vector of additive marker effects, and ϵ the vector of random errors. In this study, the marker effects in the GP model were estimated by using a ridge-regression best linear unbiased prediction (rrBLUP) model (Endelman, 2011) and Bayes A model (Pérez and De Los Campos, 2014) based on the above-described 988 clones genotyped at 25,602 markers. We incorporated singular value decomposition (SVD) to reduce the dimensions of the marker matrix to facilitate the computation in the GP model (Meuwissen et al., 2017). Subsequently, 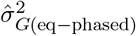 and 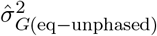 for each bi-parental cross was calculated, respectively.

In theory, the covariance between two unlinked loci is 0 in the case of c = 0.5. This would correspond to a ideal (infinite) family size in Hardy Weinberg equilibrium. However, in breeding population of a finite or small size under selection or genetic drift, the covariance between two unlinked loci could differ from 0. Therefore, we quantified the total covariance of all pairs of unlinked loci in our empirical data as:

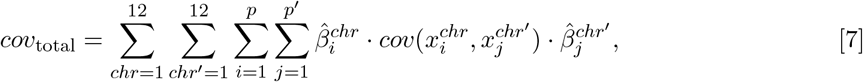

where 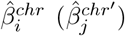 was the estimated effect of the i(j)^th^ marker at the chr(chr^′^)^th^ chromosome obtained from the GP model [6], 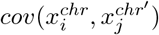 the covariance between the marker i and the marker j among the empirical progenies where the two markers were located on two different chromosomes (i.e., chr and chr^′^), and p and p^′^ the number of markers at the chr^th^ and chr^′th^ chromosomes. The cov_total_ needs to be added to 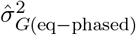 in order to obtain the predicted segregation variance under finite family sizes, which was denoted as 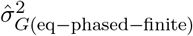, i.e., 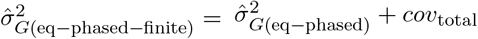.

### Evaluation of prediction performance

The segregation variance estimated from the empirical phenotypic dataset via a LMM 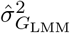 and the one from algebraic derivations 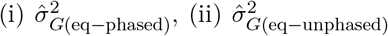 and 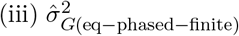 were compared by using PCC and CV[RMSD] among the 15 families with a family size > 10, where CV[RMSD] is the coefficient of variation of the RMSD to enable comparing all assessed traits with different units, and calculated as 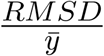, in which 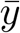 was the average of adjusted entry means among the 988 clones.

## RESULTS

### Simulation study

Our results showed that the PCC between the variance estimated using the simulated progenies 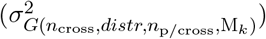 and the one estimated from the derived formula with the phased information 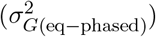 increased with an increase in family size (Figures 2a and 2b). At the same time, the RMSD between the two variances decreased with increasing family size (Figures 3a and 3b). When the family size was 5000, the PCC reached 0.96 and RMSD was smaller than 4.1 for all phased scenarios. Therefore, our derived formula with phased parental haplotypes and a genetic map can be used to precisely predict the segregation variance.

**Figure 2:**
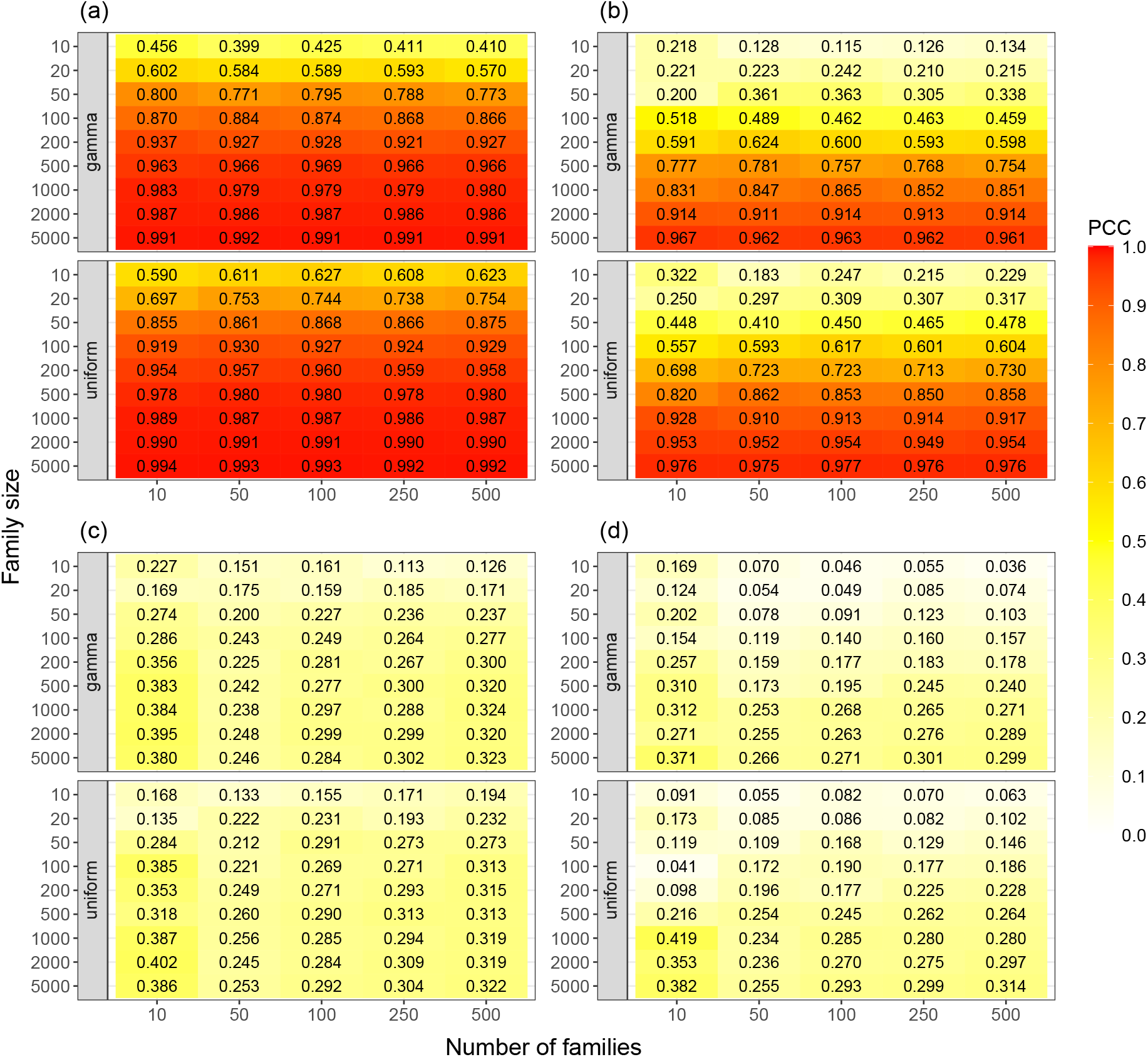
The Pearson correlation coefficient (PCC) between (a) 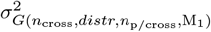 and 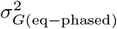; (b) 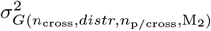 and 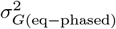; (c) 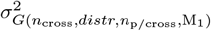 and 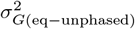; (d) 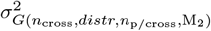 and 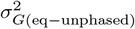 across the 30 runs. 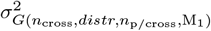: the segregation variance obtained from the summation of sample variance of true genetic values among all progenies per chromosome; 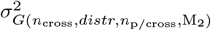: the segrega-tion variance obtained from the sample variance of true genetic values among the simulated progenies and chromo-somes; 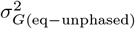: the segregation variance estimated by the derived formula with unphased parental haplotypes; 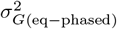: the segregation variance estimated by the derived formula with phased parental haplotypes; n_cross_: the number of families; distr: the distribution of the family sizes across the families: uniform or gamma distribution; and n_p/cross_: the family size.

**Figure 3:**
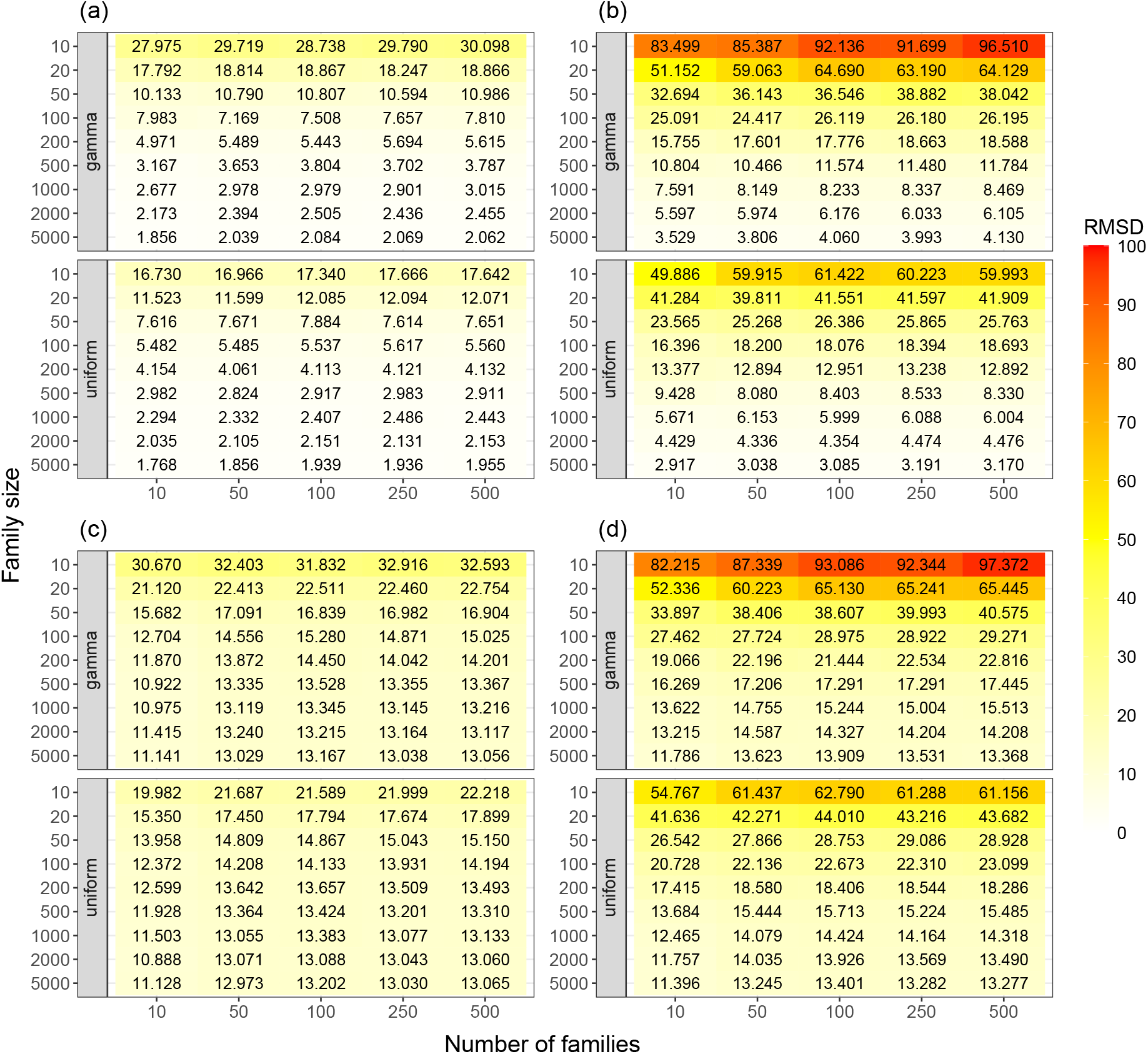
The root mean square deviation (RMSD) between (a) 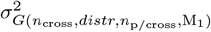 and 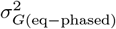; (b) 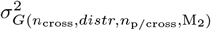 and 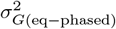; (c) 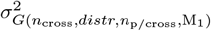 and 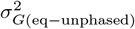; (d) 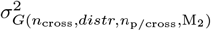 and 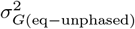 across the 30 runs. 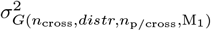: the segregation variance obtained from the summation of sample variance of true genetic values among all progenies per chromosome; 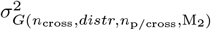: the segrega-tion variance obtained from the sample variance of true genetic values among the simulated progenies and chromo-somes; 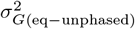: the segregation variance estimated by the derived formula with unphased parental haplotypes; 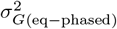: the segregation variance estimated by the derived formula with phased parental haplotypes; n_cross_: the number of families; distr: the distribution of the family sizes across the families: uniform or gamma distribution; and n_p/cross_: the family size.

**Figure 4:**
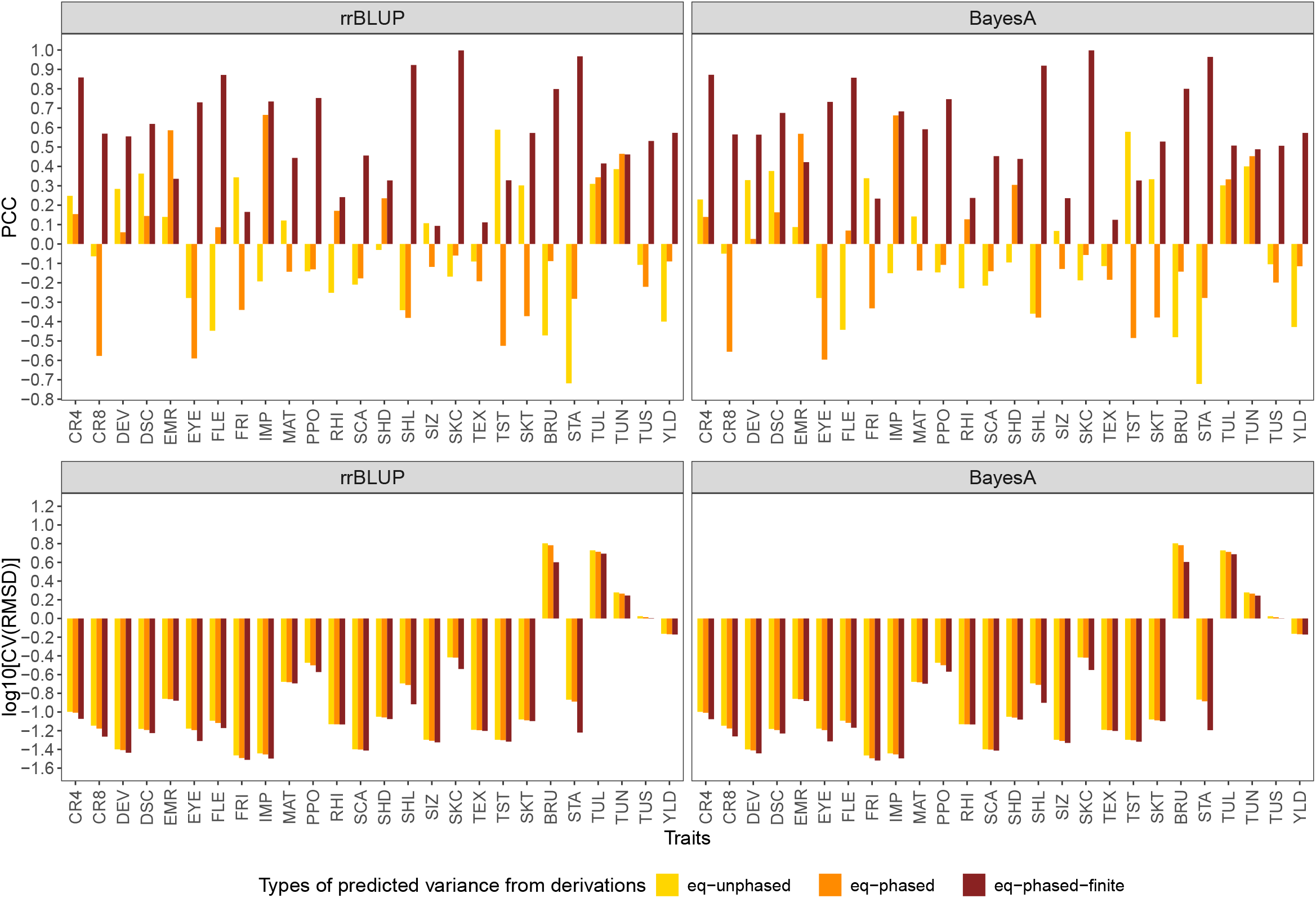
Pearson correlation coefficients (PCC, upper) and log10 of the coefficient of variation of the root mean square deviations (CV[RMSD], bottom) between segregation variances of 15 families estimated from a linear mixed model 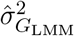 in empirical data and the ones estimated by the derivations incorporating estimated marker effects from rr-BLUP and BayesA methods. The segregation variances estimated from the derivations include (i) 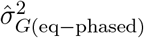: with phased parental haplotypes, (ii) 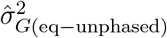: with unphased parental haplotypes, and (iii) 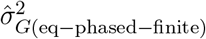: under a finite family size.

In accordance to the above-mentioned comparison, the PCC between 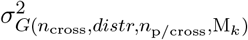 and the variance estimated using the derived formula without phased parental haplotypes 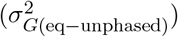 increased (Figures 2c and 2d) and their RMSD decreased with an increase in family size (Figures 3c and 3d). However, the PCC was much lower and the RMSD much higher between 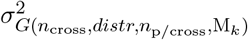 and 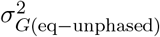 than the one between 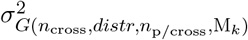 and 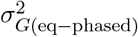. Moreover, the PCC between 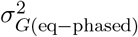 and 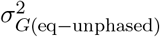 was relatively low (~ 0.3) and their RMSD high (~ 14; Table S3), independently of the number of families.

In further analyses, the PCC and RMSD between 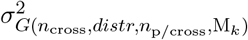 and 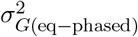 were used to assess how different parameters of experimental datasets affect the performance of predicting segregation variance. The considered parameters were the combination of (i) the different family size per segregating population, (ii) the distribution of the family size (i.e., uniform and gamma distribution), and (iii) the number of families in a breeding program. An increased PCC and a decreased RMSD were observed with an increase in family sizes (Figures 2 and 3). Furthermore, the segregation variance excluding the covariance between unlinked loci (i.e., 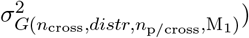 had a higher PCC and a lower RMSD than the one including the covariance of the unlinked loci (i.e., 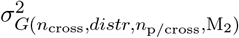 Figure 2 and Figure 3). The differences in PCC and RMSD between the two evaluated methods without/with considering the unlinked loci (M_1_ and M_2_) were less pronounced when the family size increased, and were minimal in case of very high family size.

A higher PCC (Figure 2) and a lower RMSD (Figure 3) between 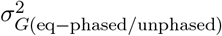 and 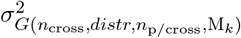 were observed for the uniform distribution of the fam-ily size than for the gamma distribution. However, the difference in PCC and RMSD between the two distributions became small as the average family size increased. Lastly, the number of families did not affect the PCC and RMSD between 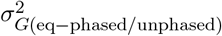 and 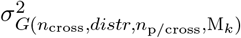.

### Prediction of segregation variance of experimental families

To assess the application of our algebraic derivations in real breeding situations, we compared the variance estimated by the derived formulae incorporating the marker ef-fects estimated in empirical potato populations (i.e., 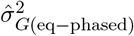 and 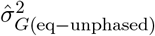) and the segregation variance estimated by the LMM from empirical data 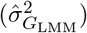. The results of all comparisons were independent of the methods used for marker estimation, i.e., rrBLUP and Bayes A (Figure 4). Therefore, we present in the following the results from rrBLUP only.

The PCC between the empirical variance 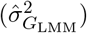 and the one estimated from the derived formulae differed largely across all the assessed traits, and ranged from −0.718 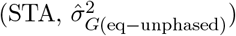 to 0.665 (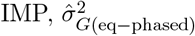 Figure 4). On average across all the assessed traits, the PCCs were negative between 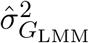 and (1) 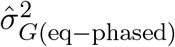 (−0.028) and (2) 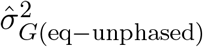 (−0.053). In detail, only around half of the traits had better PCC with 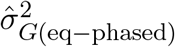 compared to the one with 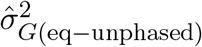. On the contrary, the CV[RMSD] with 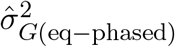 was lower for all traits compared to the one with 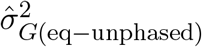 (Figure 4). The degree of this reduction depended on the traits and ranged from 0.19% (RHI) to 6.99% (CR8).

Because the family size of our empirical data were small and a higher covariance between unlinked loci was observed in small families in our simulations, we considered an additional variance, 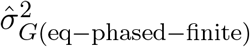 by adding the covariance of unlinked loci to 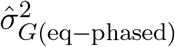. This led to considerable increases of the PCC ranging from 10.39% (IMP) to 1776.29% (SKC) (Figure 4). Only for the traits TUN and EMR, no increase was observed. These PCCs reached an average value of 0.56 across the traits. Nine out of the 26 traits had PCC values higher than 0.7. Furthermore, the CV[RMSD] of 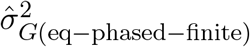 decreased dramatically compared to the CV[RMSD] of 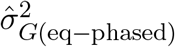, where the decrease ranged from 0.31% (RHI) to 53.32% (STA). This reduction of CV[RMSD] was more pronounced than the reduction observed from the CV[RMSD] of 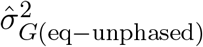 to CV[RMSD] of 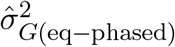 for most traits. Thus, the covariance of unlinked loci contributed a large proportion to the empirical segregation variance.

## DISCUSSION

Algebraic derivations to predict segregation variance by incorporating parental haplotype information, recombination rates, and estimated marker effects have been developed for diploid species in recent years (Bonk et al., 2016; Lehermeier et al., 2017b; Osthushenrich et al., 2017; Allier et al., 2019; Wolfe et al., 2021; Niehoff et al., 2024). However, these approaches are not suitable for autotetraploid crops. Therefore, in this study, we first focused on developing algebraic derivations for the segregation variance in autotetraploid crops with a heterozygous genome, and compared their performance to the segregation variance estimated in simulated progenies. Furthermore, we investigated how different parameters affect the accuracy of estimating segregation variance via simulations. Lastly, we demonstrated the potential of the derived formula for predicting segregation variance in potato breeding programs.

### The validity of the derived formulae

The segregation variance obtained by our algebraic derivation with phased parental haplotypes was highly correlated (0.99) with the segregation variance from simulated progenies with a huge family size (Figures 2a and b). Similar results were also observed between the derived formula and simulated progenies in diploid systems (Osthushenrich et al., 2017; Neyhart et al., 2019), even though with PopVar (Mohammadi et al., 2015) a different simulator was used in their studies compared to AlphaSimR (Gaynor et al., 2021) used in our study.

The observation of a high concordance of the algebraic derivation as well as the computer simulations was expected as both approaches rely on the same underlying information: recombination rates, phased parental haplotypes, and marker effects.

Furthermore, in this study, no double reduction was considered for both approaches. However, a small difference in segregation variance between the two approaches was observed (Figures 3a and b), which can be attributed to two main factors. First, the AlphaSimR simulator uses a stochastic process, whereas the derivation developed in our study is based on deterministic algebra. Second, AlphaSimR assumes crossover interferences in its default setting (Gaynor et al., 2021). In contrast, our algebraic derivation assumes no crossover interference (i.e., using Haldane mapping function; Haldane (1919)). A priori, we could also consider Kosambi mapping function (Kosambi, 1943) in the formula. This would eliminate the second factor. Nevertheless, with a correlation higher than 0.99 and a low error, our algebraic derivation predicts the segregation variance as precisely as simulating software.

Both approaches, the computer simulations and the algebraic derivation, are based on the assumption of the availability of phased parental haplotypes. However, due to the complexity of the genome of autotetraploids, the methodologies for their haplotype phasing remain a costly and time-consuming process (e.g., Sun et al. 2022, Sun et al. 2025). Further, the simulation approach cannot be used directly to predict segregation variance if the phased parental hapltypes are not available. In contrast, our algebraic derivation 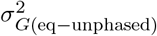, which assumes the absence of LD (i.e., covariance between loci is equal to 0), is an alternative to predict segregation variance in such cases. When correlated with the simulation results which assume that phased parental haplotypes are available, a lower PCC and higher RMSD were observed for the unphased derivation 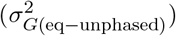 compared to the phased one 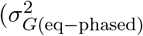; Figures 2c and 2d, Figures 3c and 3d). The same trend of low and for some extreme cases even negative PCC when correlating derivation results with the ones obtained in the empirical data was observed for both the phased and the unphased derivations (Figure 4). These observations can be explained by the fact that the LD between loci contributes to the genetic variance (Lehermeier et al., 2017a). Nevertheless, 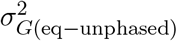 is able to catch part of the segregation variance, con-trary to the simulation approach which is unusable in the absence of phased parental haplotypes.

With regard to computational time, our derived formula in its current implementation in R does not perform quicker the prediction of segregation variance than the simulations, which is in contradiction to former studies which developed derivations for diploid systems (Lehermeier et al., 2017b; Neyhart et al., 2019). The former studies focused on inbred line breeding programs, which require the simulation of huge family sizes in each advanced generation (e.g., F2 to F8) to allow the same conclusions as the algebraic derivation. In contrast, our derivations are designed for autotetraploid clonal breeding programs, implying that each cross from two parents only needs to generate its offspring once, as clonal genotypes are maintained vegetatively in the further generations. However, the computational time to calculate segregation variance based on our algebraic derivation could be reduced by implementing parallel calculations of chromosomes and crosses, or programming in more efficient languages (e.g., C++, Java, etc).

In short, our derivations can be applied to situations with either available or unavailable phased parental haplotypes and avoid the need of simulation tools to create a huge family size, which is presumably the easier way to exploit information about segregation variance in practical breeding programs.

### The factors affecting the con/discordance between segregation variance from algebraic derivation and the one based on experimental data

The concordance between segregation variance predicted by our algebraic derivation and the one observed in our empirical data measured as PCC was low and for some traits even negative (Figure 4). Therefore, we investigated the role of several potential factors influencing the accuracy of segregation variance estimation.

### Bias in estimation of segregation variance based on experimental data

Our simulation results showed an increasing PCC and decreasing RMSD between the segregation variance estimated by the derivation and the one estimated by simulated progenies (Figures 2 and 3) with an increasing family size. Moreover, PCC and RMSD reached a plateau at around 1000 progenies (Suppl. Figure S3). This observation can be expected from a statistical perspective, as a larger family size results in a better estimation of the population variance by the sample variance, and is also in accordance with the findings of Oget-Ebrad et al. (2024). However, the family size used to assess the performance of segregation variance prediction in our empirical data was considerably lower and ranged only from 11 to 23. This could explain why low and even negative PCCs were observed between segregation variance calculated from the derived formulae and the linear mixed model 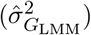 in our empirical data across the 26 examined traits (Figure 4).

Further, in small families as such that can typically be found in breeding programs, genetic drift is a major force generating significant LD between linked but also unlinked loci (Stich et al., 2007), thereby impacting the segregation variance in addition to the LD created by selection. This was confirmed by the observation that the correlation between segregation variance calculated from our derivations and simulations ignoring the LD between unliked loci (M_1_) was always higher than the one including the unliked LD (M_2_; Figures 2 and 3). Further, the difference between the correlations using methods excluding or including the covariance between unlinked loci (M_1_ versus M_2_) increased as the family size decreased. One reason for the difference between M_1_ and M_2_ could be the higher LD level in families of small size (Waples, 2006). To confirm this, we estimated the average R^2^ across all pairs of unlinked loci in the simulated progenies. Our results showed that the R^2^ decreased as the family size increased (Suppl. Figure S4). A similar pattern was observed in our analyses of the empirical dataset. In detail, the PCCs across the assessed traits dramatically increased and RMSD reduced after incorporating the covariance between the two unlinked loci in case of small family sizes (Figure 4). Therefore, we concluded that family size strongly affects the predictive performance of segregation variance, because a small family size results in an increased covariance between unlinked loci.

The distribution of the family sizes across segregating populations was different between the breeding programs (e.g., Suppl. Figure S2). This factor could also affect the prediction performance of segregation variance and, to our knowledge, has not been evaluated yet. Therefore, we compared the segregation variance prediction for different distributions of the family size. In our simulation results, under the same average family size, the gamma distribution showed a lower PCC and a higher RMSD than the uniform distribution (Figures 2 and 3). The differences in PCC and RMSD between the two distributions were more pronounced with a decrease in the average family size. One explanation is that, when following a gamma distribution, some families are smaller than the average, which could lead to worse performance in predicting segregation variance compared to the uniform distribution with evenly sized families. In contrast, the number of families did not affect the accuracy of predicting segregation variance (Figures 2 and 3). This factor only affects statistical power, which is in line with the trend of an increased significance of the correlation with a greater number of families (Suppl. Figure S5).

### Precise estimation of segregation variance via algebraic derivation

According to equation [1], the two main components, marker effects and additive genotypic variance of the progenies directly influence the accuracy of the algebraic derivation in predicting segregation variance. Thus, to increase the accuracy of the prediction of segregation variance, accurately estimating marker effects plays an important role. Several factors, including the choice of GP models, the relatedness between training and validation set, the number of markers, and the size of the training set have been proven to influence prediction ability and therefore marker effect estimation (see review: Alemu et al. 2024). For the first factor, the choice of the GP model, only minor differences in prediction abilities were observed between the rrBLUP and Bayes A models (Suppl. Figure S6), and, thus, the performance in predicting segregation variance between the two GP models was similar. In addition, different genetic architectures and the heritabilities of the trait also affect the accuracy of estimated marker effects. This was in accordance with our results and the ones from Thelen et al. (2025, in press), in which the prediction abilities between observed and estimated genetic values across the assessed traits varied consequently (Suppl. Figure S6). A similar trend of different performance across the traits was found in our empirical analysis when correlating the segregation variance estimated from the mixed linear model with the one estimated from the derived formulae (Figure 4). This was in accordance with former studies on barley (Osthushenrich et al., 2017), cassava (Wolfe et al., 2021), soybean (Wartha and Lorenz, 2024), and wheat (Oget-Ebrad et al., 2024). In summary, a precise estimation of the marker effects is necessary to obtain reliable segregation variance estimation from the algebraic derivations.

The second component influencing the accuracy of our derivation is the additive genotypic variance of the progenies. Therefore, both the assumptions and the accuracy of the input information required for the algebraic derivation affect its precision. The first assumption made was that only bivalent form during meiosis, which implies that no double reduction occurs. The second was that the conversion between the genetic map and the recombination rate between two markers followed a diploid-like inheritance, which ignores the effect of double reduction as well (Zheng et al., 2021). However, the phenomenon of double reduction can be found in autotetraploid species (Gallais, 2003; Bourke et al., 2015), and therefore, its effect on predicting segregation variance requires further investigation.

With regard to the required inputs for the calculation of the derivation of the segregation variance, the algebraic derivation relies on accurate phased parental haplotypes and recombination rates. To infer the phased parental haplotypes in this study, we used the PolyOrigin software (Zheng et al., 2021), which recommends having at least 30 progenies to construct the parental haplotype. However, the family size across the 15 families in our study was lower than 30. Therefore, in order to increase the correlation between the segregation variance estimated using the algebraic derivation and the one obtained from the empirical data by having more reliable phased parental haplotypes, bigger families are required than used in our study. Regarding the recombination rates, those were fixed across different segregating populations in our derivations, although different segregating populations are known to have dif-ferent crossover events (Casale et al., 2022; Jiang et al., 2023). These factors could impact the precise estimation of the segregation variance.

In a quantitative genetics perspective, the genotypic variation is attributed not only to additive but also non-additive effects. Furthermore, autotetraploid potato has a highly heterozygous genome, which can be accompanied by important dominance effects for quantitative traits. The existence of dominance effects across various traits has been shown in different potato breeding materials (Endelman et al. 2018 and K. Thelen personal communication). However, only additive effects were considered in our derivation. This could lead to imprecise estimation of the segregation variance, especially for a trait with strong non-additive effects. Thus, incorporating non-additive effects, such as dominance effects, to predict segregation variance is relevant but requires further research.

### Applying segregation variance prediction in clone breeding programs

Optimizing the selection of superior crosses to improve the genetic gain is one of the most important steps in practical breeding programs. According to the literature (Schnell and Utz, 1975; Zhong and Jannink, 2007), not only the progeny mean but also the segregation variance — and their respective variances — play a pivotal role in determining potential crosses. Several possibilities are available for predicting the mean (e.g., Endelman, 2025), but not the segregation variance of autotetraploid species. Therefore, developing an accurate estimation of the segregation variance becomes necessary for breeding programs. In this study, we derived novel algebraic derivations applied to autotetraploid species to predict segregation variance. These derivations incorporating the marker effects from GP models can be used to predict segregation variance and therefore pre-select the potential parental combinations for the crossing plan. Furthermore, the derivations can help to overcome limitation in the field, where it is impossible for the breeders to grow all families with a large family size. Knowledge about segregation variances might lead to improved resource allocation, as breeding companies can select between segregating populations with equally high means (Mohammadi et al., 2015). Subsequently, breeders could classify their potential crosses according to the expected segregation variance using the presented derivation and, thus, decide to produce huge family only for promising crosses with high progeny mean and high segregation variance. Therefore, the proposed derivations promise to be a precise alternative to simulations and empirical populations to help breeders in selecting the crosses and their family sizes considering segregation variance. By this, the proposed derivations can be used for optimizing the breeding programs, as segregation variance is a parameter needed for their simulation when trying to increase the long-term genetic gain and monitor diversity in autotetraploid clonal breeding programs (Wu et al., 2025).

## ACKNOWLEDGMENTS

This study was funded by the Federal Ministry of Food and Agriculture/Fachagentur Nachwachsende Rohstoffe (grantID 22011818 – PotatoTools and grantID 2222NR078A – PotatoPredict). We thank Dr. Chaozhi Zheng for helping to use the PolyOrigin software. This work was supported by the de.NBI Cloud within the German Network for Bioinformatics Infrastructure (de.NBI) and ELIXIR-DE (Forschungszentrum Jülich and W-de.NBI-001, W-de.NBI-004, W-de.NBI-008, W-de.NBI-010, W-de.NBI-013, W-de.NBI-014, W-de.NBI-016, W-de.NBI-022). Computational infrastructure was also provided by the central Information Technology at IPK Leibniz Institute. The funders had no influence on study design, the collection, analysis and interpretation of data, the writing of the manuscript, and the decision to submit the manuscript for publication.

## SUPPORTING INFORMATION

Additional supporting information can be found online in the Supporting Information section at the end of this article.

## DATA AVAILABILITY

The datasets generated or analyzed during this study along with the R scripts will be made available upon acceptance.

## AUTHOR CONTRIBUTIONS

Po-Ya Wu: conceptualization, methodology for the derivation, simulations, data analysis, visualization, and writing; Kathrin Thelen: empirical data analysis and writing; Benjamin Stich: conceptualization, funding acquisition, project administration, and writing; Stefanie Hartje, Katja Muders, and Vanessa Prigge: funding acquisition, resources, and review; and Delphine Van Inghelandt: conceptualization, methodology for the derivation, funding acquisition, project administration, supervision, and writing. All authors read and approved the final manuscript.

## CONFLICT OF INTEREST

Vanessa Prigge is an employee of Saka Pflanzenzucht GmbH & Co. KG. Katja Muders is an employee of NORIKA GmbH. Stefanie Hartje is an employee of EUROPLANT Innovation GmbH & Co. KG. The authors declare no conflict of interest.

## SUPPLEMENTARY MATERIAL

Table S1: The complete 1296 possible genotypes in a F1 family of a bi-parental cross and their corresponding probabilities, depending on the parental gametes and their probability. c is the recombination rate between loci A and B.

File: all_genotype_1296_combination_Suppl.xlsx

**Table S2:** Description of the 15 environments (i.e., year-location combinations) that were used in the experiment and their respective properties. The breeding companies are Europlant (EUROPLANT Innovation GmbH & Co. KG), Norika (NORIKA GmbH), and SaKa (SaKa Pflanzenzucht GmbH & Co. KG).

| Environment | No. of<br>entries | No. of<br>families | No. of<br>blocks | No. of<br>plants per plot |
| --- | --- | --- | --- | --- |
| Europlant 2019 Kaltenberg | 299 | 46 | 4 | 10 |
| Europlant 2020 Kaltenberg | 297 | 47 | 4 | 16 |
| Europlant 2020 Böhlendorf | 287 | 46 | 2 | 16 |
| Europlant 2021 Kaltenberg | 300 | 47 | 4 | 16 |
| Europlant 2021 Böhlendorf | 300 | 47 | 1 | 16 |
| Norika 2019 Groß Lüsewitz | 300 | 17 | 2 | 9 |
| Norika 2020 Groß Lüsewitz | 300 | 17 | 4 | 18 |
| Norika 2020 Mehringen | 300 | 17 | 3 | 20 |
| Norika 2021 Groß Lüsewitz | 297 | 17 | 4 | 18 |
| Norika 2021 Mehringen | 300 | 17 | 2 | 20 |
| Saka 2019 Windeby | 458 | 107 | 8 | 10 |
| Saka 2020 Windeby | 387 | 99 | 8 | 16 |
| Saka 2020 Gransebieth | 387 | 99 | 8 | 16 |
| Saka 2021 Windeby | 387 | 99 | 8 | 16 |
| Saka 2021 Gransebieth | 387 | 99 | 8 | 16 |

**Table S3:** Medians of Pearson correlation coefficient (PCC) and root mean square deviations (RMSD) between segregation variance predicted from the algebraic derivations with 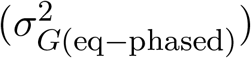 and without 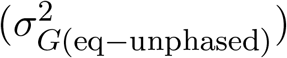 phased parental haplotypes using different number of families (n_cross_ = 10, 50, 100, 250, and 500) among 30 simulation runs.

| | | $n_{\text{cross}}$ | | | | |
| --- | --- | --- | --- | --- | --- | --- |
|  |  | 10 | 50 | 100 | 250 | 500 |
| PCC |  | 0.377 | 0.217 | 0.269 | 0.283 | 0.311 |
| RMSD |  | 11.808 | 14.045 | 14.171 | 13.937 | 14.103 |

**Figure S1:**
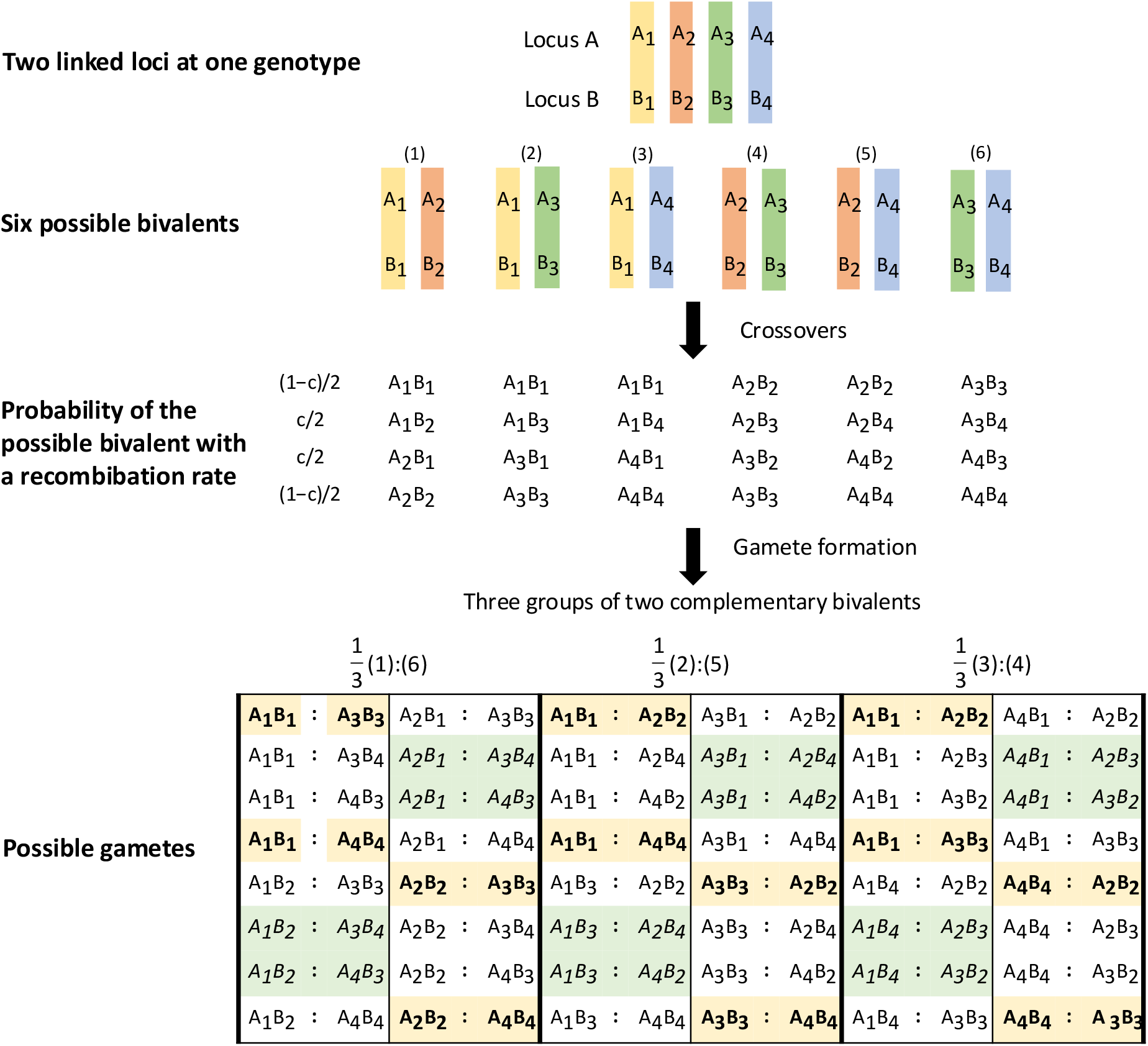
Formation of the gametes from a autotetraploid genotype (considering only random bivalents during meiosis). A genotype at two linked loci, A and B, with a frequency of recombination c across four homologs are expressed A_1_B_1_/A_2_B_2_/A_3_B_3_/A_4_B_4_. Ignoring linkage phases (coupling and repulsion), there are 16 types of gametes for each pair of groups but in total there are 36 types of gametes instead of 48. This is because both gametes with parental associations (in bold) and those with two recombined chromosomes (in underline) are present twice. This figure is revised from Figure 1.4 in Gallais (2003).

**Figure S2:**
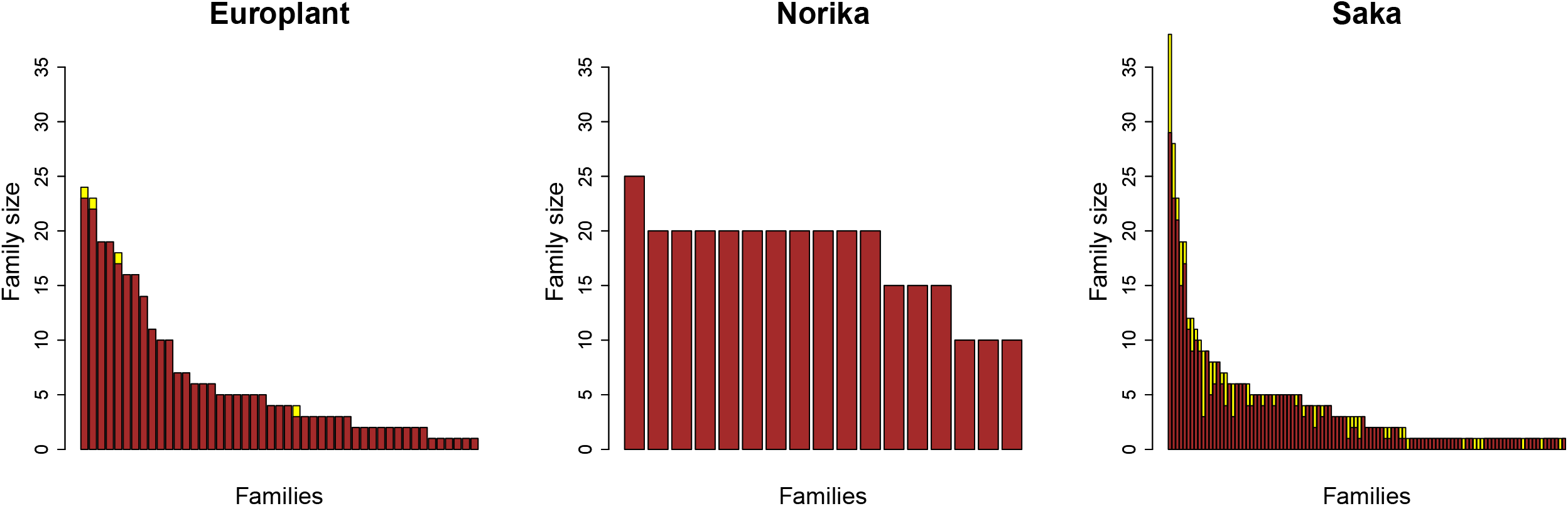
The distribution of the family size of the 171 families across the three breeding companies: Europlant (EUROPLANT Innovation GmbH & Co. KG), Norika (NORIKA GmbH), and Saka (SaKa Pflanzenzucht GmbH & Co. KG) at A clone stage, respectively. The bar filled in yellow represented the 171 families among the 1058 phenotyped entries and the bar in brown the 163 families among the 980 genotyped entries.

**Figure S3:**
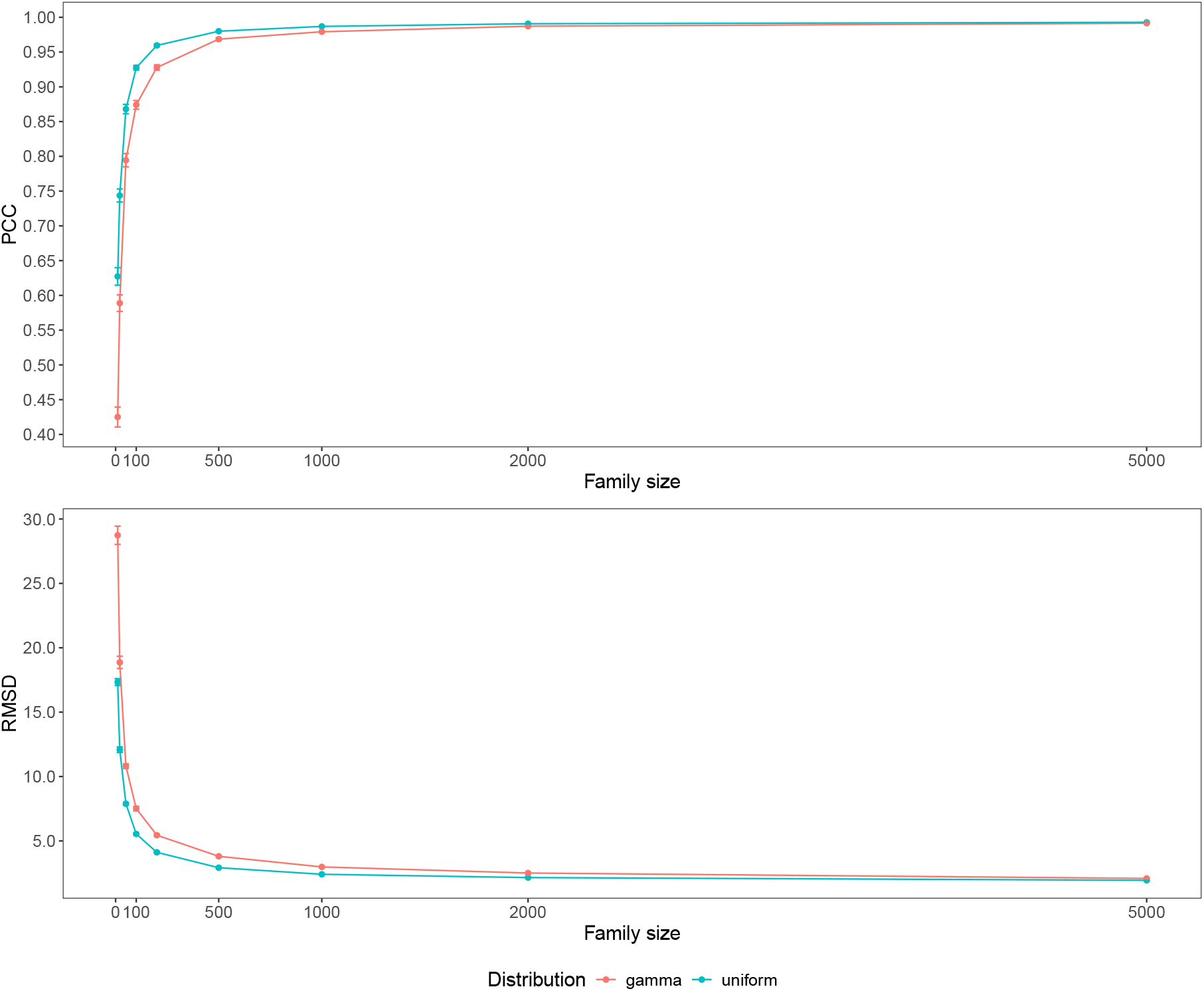
Plot of median of Pearson correlation coefficient (PCC, upper) and root mean square deviation (RMSD, bottom) across the 30 runs against different family sizes. The error bar is the standard error of PCC and RMSD across the 30 runs. The PCC and RMSD were calculated between 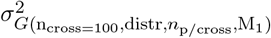 and 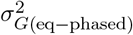. 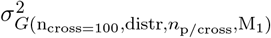: the segregation variance obtained from the summation of sample variance per chromosome; 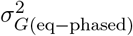: the segregation variance estimated by the derived formula with phased parental haplotypes; n_cross_: the number of families (100 here); distr: the distribution of the families size across families: uniform or gamma distribution; and n_p/cross_: the family size.

**Figure S4:**
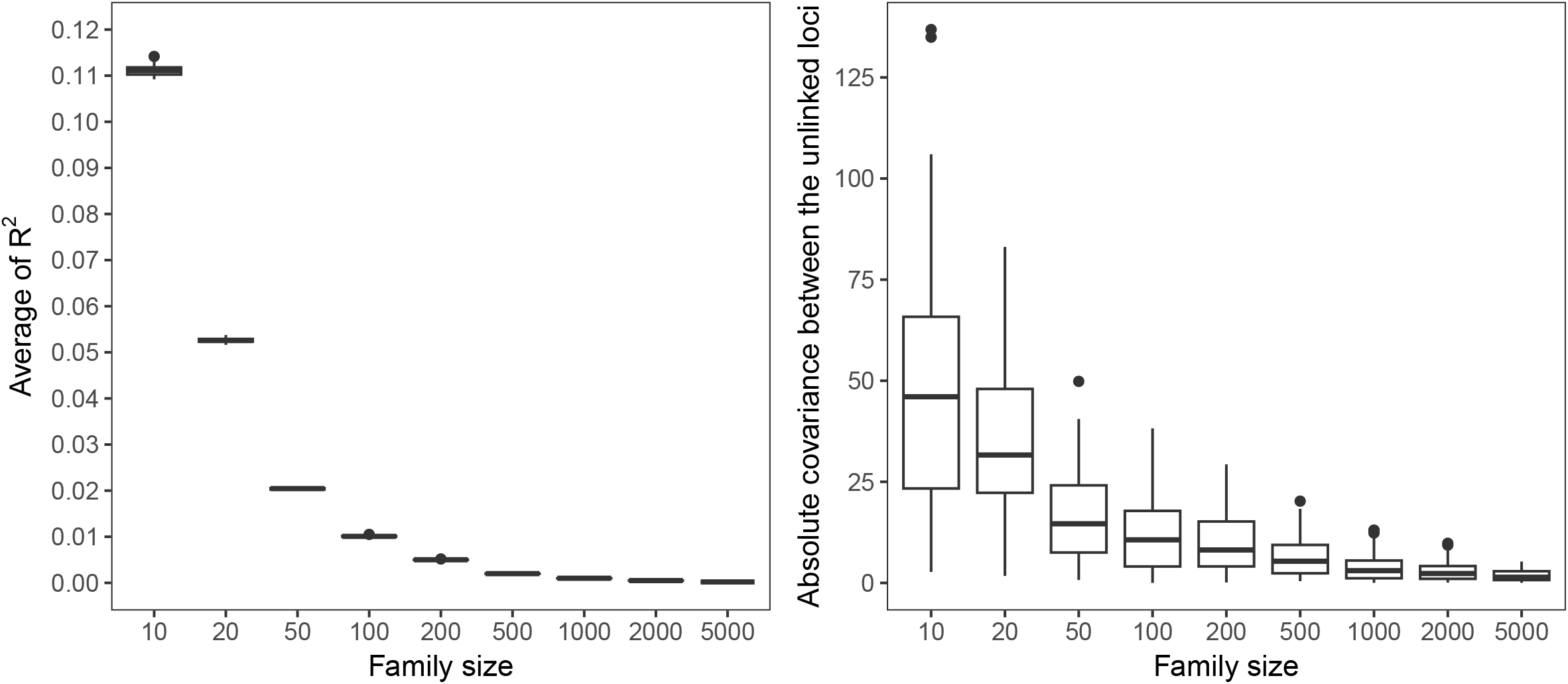
Boxplot of the average of R^2^ among all pairwise unlinked loci (left), and the absolute total covariance between the unlinked loci (right), against different family sizes across 50 families. R^2^ was calculated as the square of the Pearson correlation coefficient between two unlinked loci among the simulated progenies.

**Figure S5:**
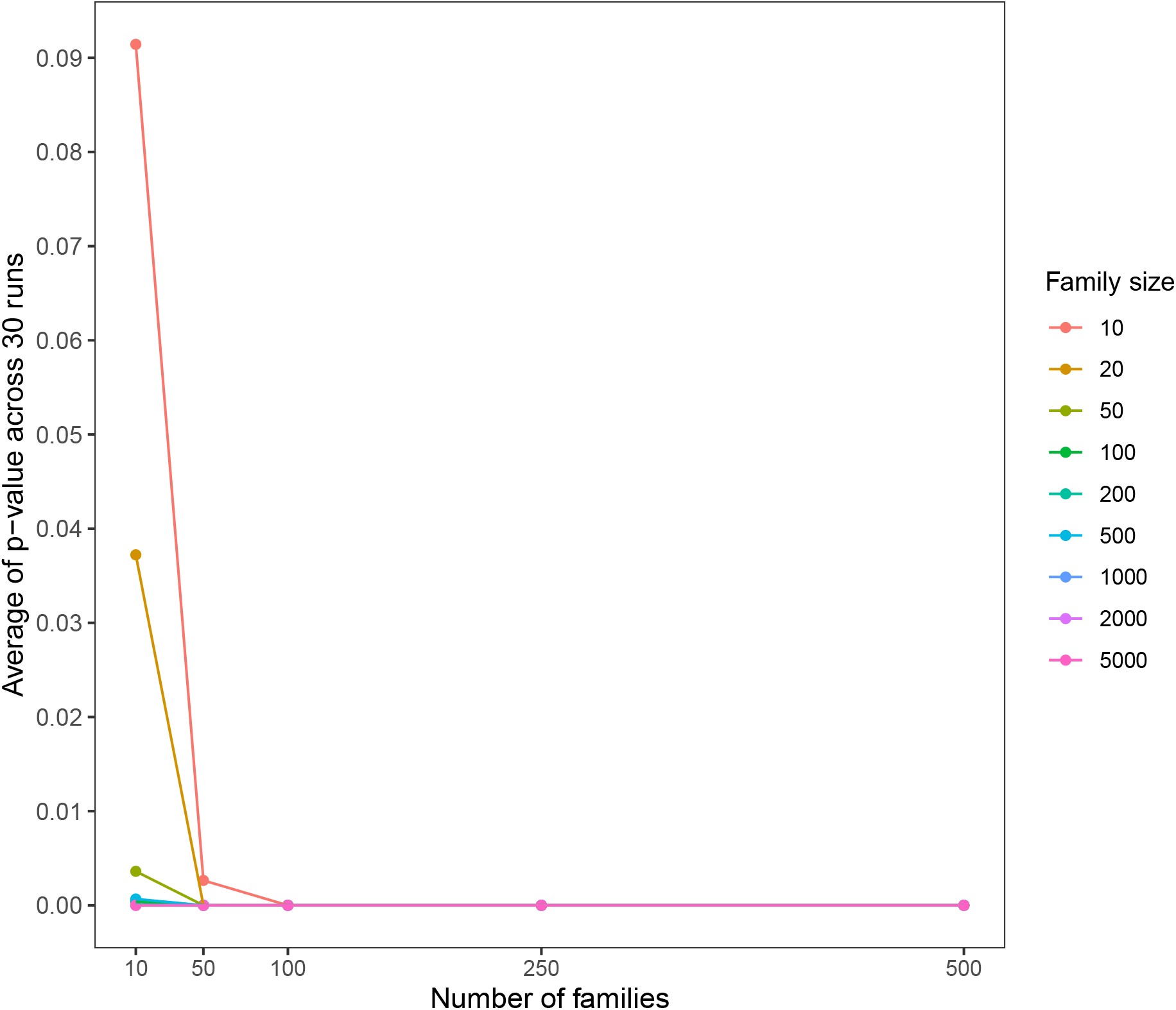
The average of the p−value of the Pearson correlation coefficient between the segregation variance from the derived formula 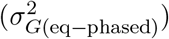 and the one estimated from simulated progenies using different numbers of families and family sizes and ignoring the covariance between the unlinked loci 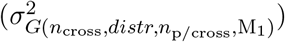 among 30 simulation runs.

**Figure S6:**
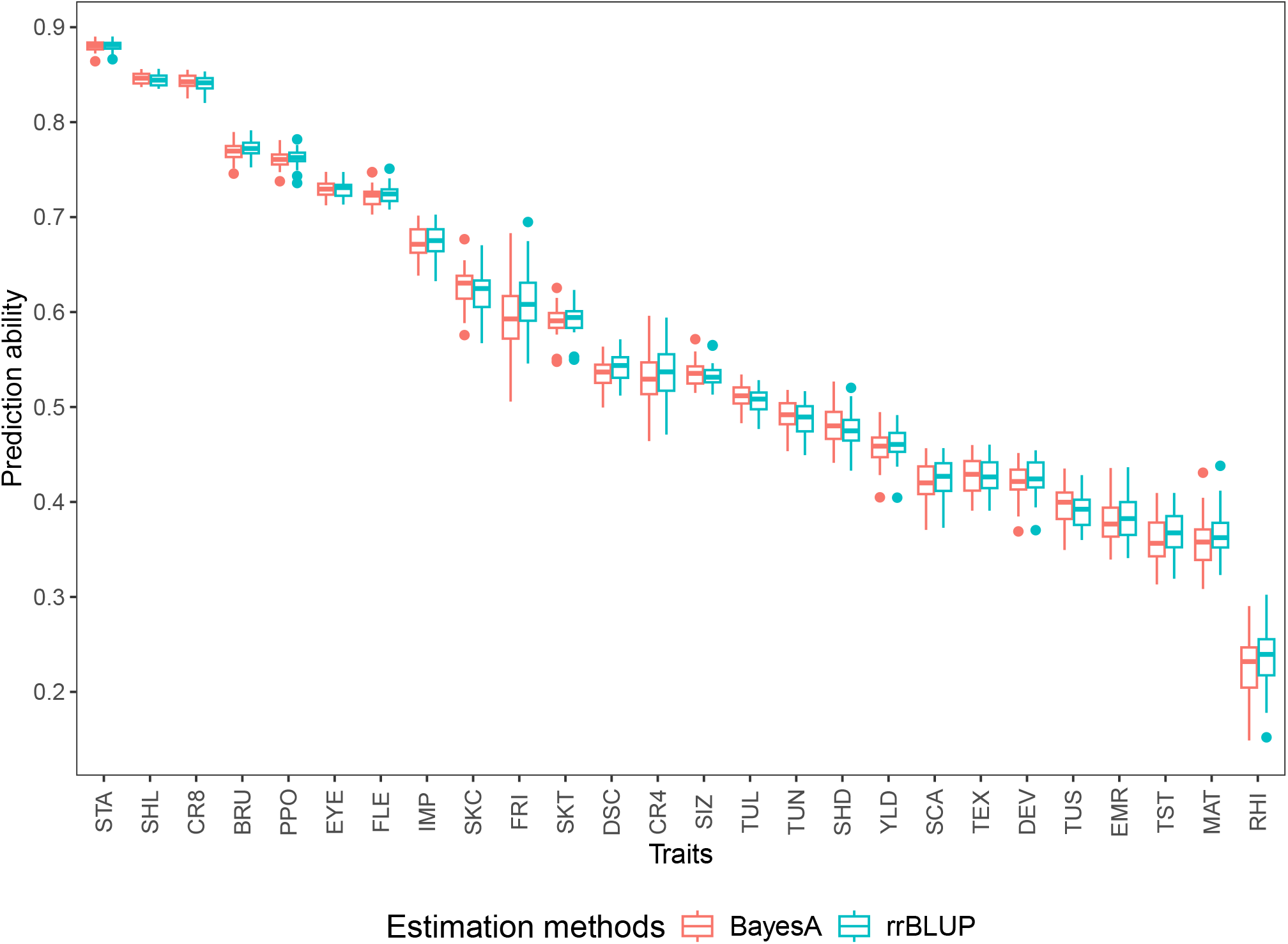
Boxplots of prediction abilities for all the assessed traits based on 988 clones and 25602 markers using rrBLUP and Bayes A estimations across 30 five-fold cross-validation (CV) runs. The prediction ability was calculated as the median of the correlation between the adjusted entry means and the estimated genetic values from the genomic prediction model among the five-fold CV within each replicate.

